# Discovery of a Gut Bacterial Metabolic Pathway that Drives α-Synuclein Aggregation and Neurodegeneration

**DOI:** 10.1101/2022.06.08.495350

**Authors:** Lizett Ortiz de Ora, Kylie S. Uyeda, Elizabeth Bess

## Abstract

Parkinson’s disease (PD) etiology is associated with aggregation and accumulation of α-synuclein (α- syn) proteins in midbrain dopaminergic neurons. Emerging evidence suggests that in certain subtypes of PD, α-syn aggregates originate in the gut and subsequently spread to the brain. However, the mechanisms that instigate α-syn aggregation in the gut have remained elusive. In the brain, the aggregation of α-syn is induced by oxidized dopamine. Such a mechanism has not been explored in the gastrointestinal (GI) tract, a niche harboring 46% of the body’s dopamine reservoirs. Here, we report that gut bacteria *Enterobacteriaceae* induce α-syn aggregation. More specifically, our *in vitro* data indicate that respiration of nitrate by *Escherichia coli* K-12 yields nitrite, a potent oxidizing agent that creates an oxidizing redox potential in the bacterial environment. In these conditions, Fe^2+^ was oxidized to Fe^3+^, enabling formation of dopamine-derived quinones and α-syn aggregates. Exposing nitrite, but not nitrate, to enteroendocrine STC-1 cells induced aggregation of α-syn that is natively expressed in these cells, which line the intestinal tract. Finally, we examined the *in vivo* relevance of bacterial nitrate respiration to the formation of α-syn aggregates using *Caenorhabditis elegans* models of PD. We discovered that nematodes exposed to nitrate-reducing *E. coli* K-12 displayed significantly enhanced neurodegeneration as compared to an *E. coli* K-12 mutant that could not respire nitrate. This neurodegenerative effect was absent when α-syn was mutated to prevent interactions with dopamine-derived quinones. Taken together, our findings indicate that gut bacterial nitrate reduction may be critical to initiating intestinal α- syn aggregation.

**Table of Contents Graphic:** 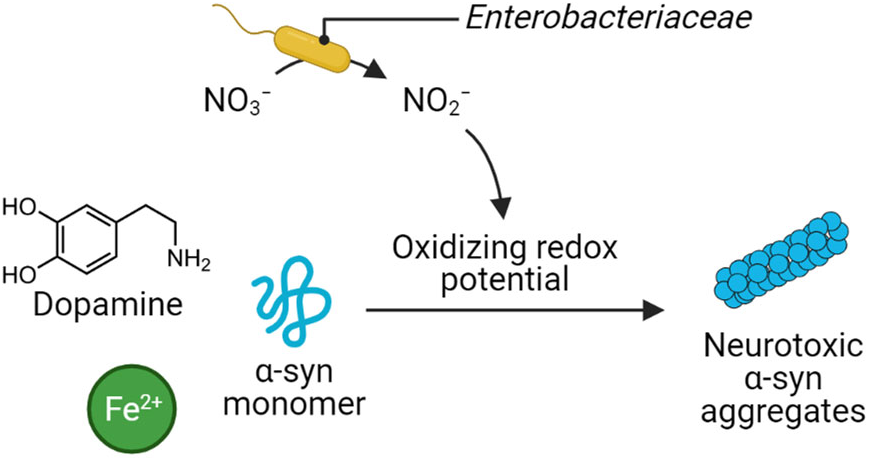

## INTRODUCTION

Although Parkinson’s disease (PD) has long been thought to originate in the brain, accumulating evidence indicates that some PD subtypes originate in the gastrointestinal (GI) tract.^1, 2^ PD is characterized by motor impairment that arises when α-synuclein (α-syn) protein aggregates accumulate in dopaminergic neurons of the substantia nigra, the brain site of motor control;^3^ however, α-syn expression is not limited to the brain. α-syn is also expressed within the mucosa of the intestinal wall by enteroendocrine cells (EECs)^4^ as well as by enteric neurons that innervate the GI tract.^5^ At least eight years prior to onset of motor symptoms in people with idiopathic PD, α-syn aggregates accumulate in GI tissue.^6^ These protein aggregates may subsequently propagate, putatively in a prion-like fashion, from the intestine to the brain via the vagus nerve that connects these organs.^7, 8^ While there is evidence that intestinal α-syn aggregates foreshadow neurodegeneration in the brain, the molecular-level mechanisms responsible for intestinal α-syn aggregation have remained poorly understood.

Several lines of evidence suggest a microbial component in the development of α-syn aggregates and progression of PD. The gut microbiota is distinct in people with PD as compared to non-diseased controls.^9–13^ This dysbiosis is often characterized by an enrichment of the facultative anaerobic *Enterobacteriaceae* bacterial family whose abundance in the gut positively correlates with the severity of motor dysfunction in people with PD.^9, 14–16^ Although it remains controversial whether gut microbiota dysbiosis is a cause or a consequence of PD pathogenesis, studies using mouse models implicate the gut microbiome in the etiology of this disorder. In germ-free mice overexpressing α-syn, GI-tract colonization using fecal samples from people with PD exacerbated motor deficits and brain pathology as compared to colonization using fecal samples from non-diseased controls.^17^ Additionally, induction of intestinal inflammation that is commonly associated with blooms in *Enterobacteriaceae* (DSS-induced colitis)^18, 19^ resulted in accumulation of α-syn in GI tracts followed by the pathogenic buildup of this protein in the brains of α-syn-overexpressing mice.^20, 21^

To identify specific gut bacterial biochemical processes that induce α-syn aggregation in the GI tract, we sought clues in characterized mechanisms of α-syn aggregation in the brain. In brain dopaminergic neurons, iron and dopamine can form a toxic pair that leads to aggregation of neural α-syn (Figure 1). Aging-related accumulation of iron in dopaminergic neurons causes oxidative stress that results in labile cytosolic ferrous iron (Fe^2+^) being oxidized to ferric iron (Fe^3+^).^22^ Cytoplasmic dopamine that is abundant in dopaminergic neurons can be readily oxidized by Fe^3+^ to highly reactive *ortho*- quinones.^23^ Dopamine-derived quinones interact with neural α-syn to cause misfolding that results in toxic α-syn oligomers.^24^

**Figure 1.**
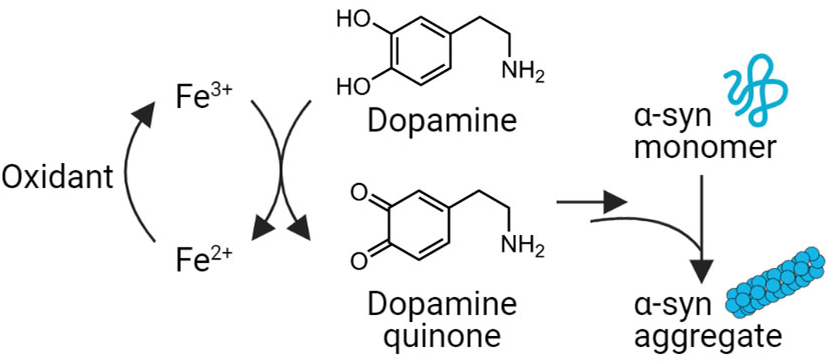
Upon oxidation of Fe^2+^ to Fe^3+^ in brain dopaminergic neurons, dopamine can be oxidized to *ortho*-quinones that cause α-syn to misfold and aggregate.

Like in the brain, the GI tract harbors dopamine and iron as well as expresses α-syn. Of the body’s dopamine pool, 46% is contained in the GI tract,^25^ and the gut microbiota is responsible for elevating the dopamine concentration in intestinal tissue.^26^ Iron, which can mediate dopamine oxidation, is present in high concentrations in the intestinal lumen (up to 25 mM), and increased dietary iron increases iron levels in intestinal cells.^27^ Although the oxidation state of labile cytosolic iron is predominantly Fe^2+^ in the non-diseased GI tract,^28^ conditions of oxidative stress increase the concentration of Fe^3+^, which could oxidize dopamine as depicted in Figure 1. With the convergence of concentrated dopamine and iron in the intestinal epithelium as well as the expression of α-syn by intestinal EECs and enteric neurons, α-syn aggregation is poised to occur in the GI tract. We sought to identify gut bacterial biochemical processes that supply the oxidant capable of inducing iron-mediated dopamine oxidation and subsequent α-syn aggregation.

Although changes in redox potential that induce oxidative stress in the GI tract are typically associated with host metabolic processes,^29–33^ gut bacteria also modulate the redox potential of their environment.^34^ Here, we describe that the ability of *Escherichia coli* (a prototypic gut bacterium of the *Enterobacteriaceae* bacterial family^18^) to create an environment with an oxidizing redox potential is a stimulus that provokes iron and dopamine to cause α-syn aggregation. We identify that nitrite, which is an oxidant generated during *Enterobacteriaceae* nitrate dissimilatory metabolism,^18, 35, 36^ stimulates a cascade of oxidation reactions that results in α-syn aggregation and neurodegeneration. Our results from *in vitro* experiments with both bacterial cultures and α-syn-expressing intestinal epithelial cells as well as *in vivo* experiments using a *Caenorhabditis elegans* model of PD suggest a novel molecular mechanism by which the gut microbiota may influence PD pathogenesis.

## RESULTS

### *E. coli* nitrate respiration creates an environment of oxidizing redox potential

Given the positive correlation between PD severity and the abundance of *Enterobacteriaceae* in the gut microbiotas of people with PD,^9^ we sought to determine whether metabolic capabilities of this bacterial family are implicated in the pathogenic aggregation of α-syn. We were particularly intrigued by *Enterobacteriaceae*’s ability to perform anaerobic nitrate respiration, which results in production of an oxidant, nitrite.^18, 37, 38^ We hypothesized that an oxidizing redox potential would be created by nitrate-respiring *Enterobacteriaceae* and, thereby, stimulate shifts in the relative abundance of labile iron from being mainly Fe^2+^ (which putatively predominates in the reducing conditions of the gut microbiota^39^) to Fe^3+^ (which could subsequently oxidize dopamine and induce α-syn misfolding and aggregation).

To test our hypothesis, we anaerobically cultured *Escherichia coli* K-12 to enable two types of metabolism: fermentation and respiration. Fermentation conditions were created by culturing *E. coli* K-12 in a minimal-nutrient medium supplemented with Fe^2+^ (500 µM) as well as glucose (20 mM) as the sole carbon source but without nitrate (media referred to as mM9_−NO3_). Conditions for nitrate respiration were generated by supplementing the same medium with nitrate (50 mM; media referred to as mM9_+NO3_). As shown in Figure 2a, supplementation with nitrate afforded a 1.6-fold increase in bacterial growth after 12 hours of incubation as measured by optical density of cultures at 600 nm (OD_600_). These findings are consistent with reported *in vivo* findings: due to nitrate respiration being more energetically lucrative than fermentative metabolism,^40^ a higher concentration of intestinal nitrate enables a bloom in *Enterobacteriaceae* abundance in the gut microbiota.^18^

**Figure 2.**
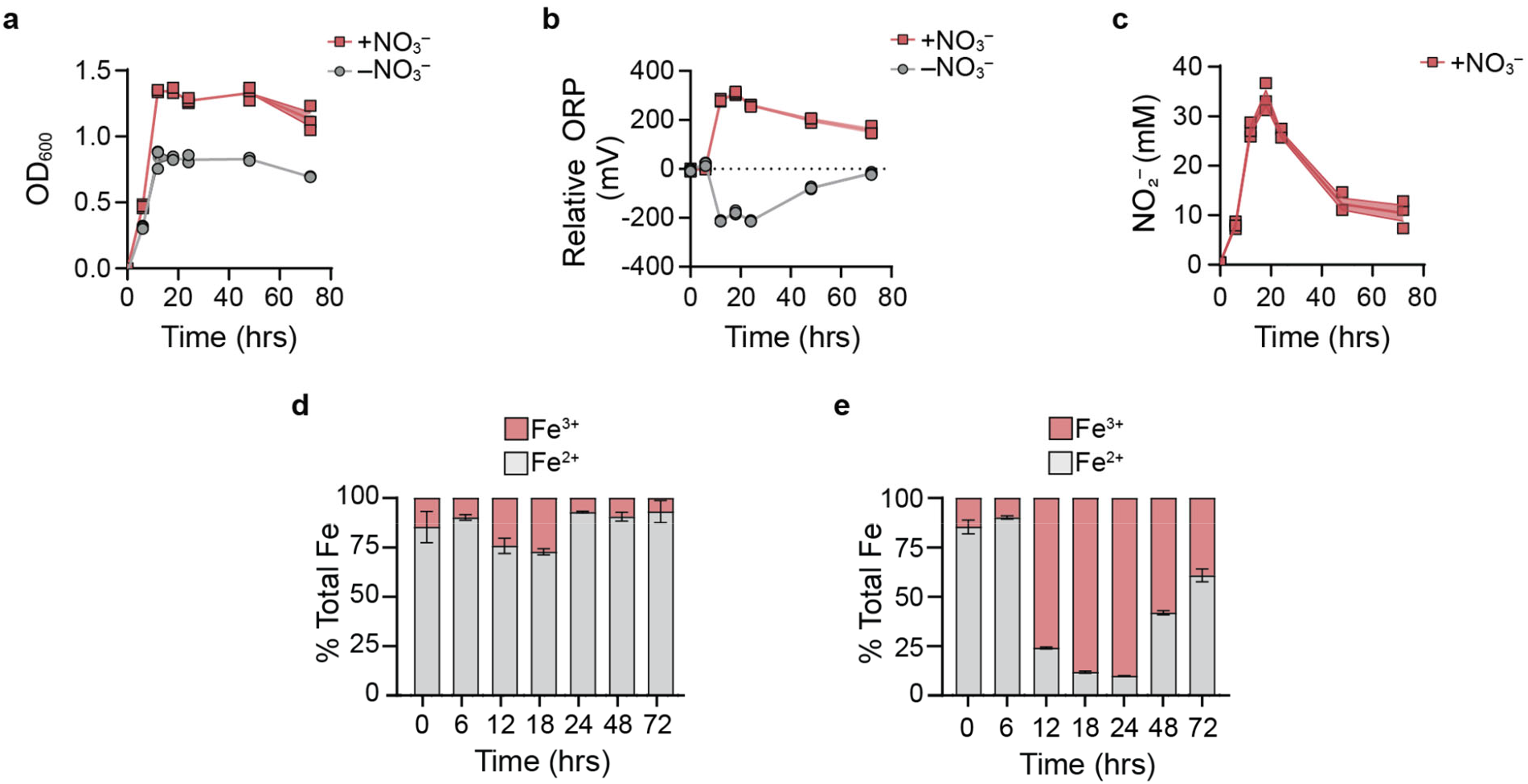
*E. coli* nitrate reduction generates an oxidant, nitrite, that creates an oxidizing redox potential in the bacterial environment and increases the relative abundance of Fe^3+^. *E. coli* K-12 was incubated in mineral media with nitrate (mM9_+NO3_) or without (mM9_−NO3_). (a) Growth was measured by optical density at 600 nm (OD_600_). (b) Oxidation-reduction potential (ORP) of bacterial cultures was measured, using an electrode, in culture media relative sterile media. (c) Nitrite was quantified in cultures supplied with nitrate using the Griess assay. (d–e) Iron speciation was measured using the ferrozine assay in bacterial cultures where (d) media lacked nitrate (mM9_−NO3_) or (e) media was supplemented with nitrate (mM9_+NO3_). n = 3 biological replicates; bars denote means ± S.E.M.

In *E. coli* K-12 cultures, we also evaluated fluctuations in redox potential as a function of bacterial metabolism. A reducing redox potential (relative to sterile controls) was observed when bacteria was cultured in mM9_−NO3_ (Figure 2b). The progressively more reducing redox potential observed over the course of exponential bacterial growth is characteristic of fermentative metabolism that yields hydrogen, a reducing agent.^41^ In contrast, bacteria cultured in mM9_+NO3_ demonstrated a steadily increasing oxidizing redox potential (relative to sterile controls) over 18 hours of incubation, reaching a maximum value of 308.17±7.62 mV (Figure 2b). Notably, increases in oxidative redox potential mirrored accumulation of nitrite in culture media (Figure 2c). Further supporting nitrite’s role as a redox-active metabolite, we observed that supplementation of sodium nitrite to sterile mM9 medium (mM9_+NO2_) afforded increases in redox potential in a concentration dependent manner (Figure S1). In contrast, uninoculated sterile media supplemented with sodium nitrate (mM9_+NO3_) showed no significant variation in the redox potential of the solution. These findings support the notion that bacterial nitrate respiration can shift the redox potential of the environment to being more oxidizing through the production of nitrite.

We next assessed whether the oxidizing redox potential afforded by nitrate-respiring conditions could shift the balance of iron speciation from favoring Fe^2+^ to favoring Fe^3+^. *E. coli* K-12 was cultured in mM9_−NO3_ or mM9_+NO3_ media, and the relative abundance of Fe^2+^ and Fe^3+^ was measured using the ferrozine assay.^42^ Under fermentation conditions that afforded a reducing redox potential (mM9_−NO3_), Fe^2+^ was the predominant iron species (Figure 2d). Conversely, in culture conditions for bacterial nitrate respiration (mM9_+NO3_), the oxidizing redox potential that corresponded to production of nitrite drove iron oxidation such that Fe^3+^ became the dominant oxidation state of iron (Figure 2e). Taken together, these findings demonstrate that the presence of nitrate enhances *E. coli* K-12 growth as well as creates an oxidizing redox potential that corresponds to nitrite production and that results in the predominance of Fe^3+^ over Fe^2+^ in culture media.

### *E. coli* nitrate respiration initiates a cascade of oxidation reactions that lead to α-syn aggregation

Next, we sought to determine whether bacterial nitrate respiration could incite the cascade of oxidation reactions that are implicated in dopamine-dependent α-syn aggregation in cerebral dopaminergic neurons but that remain unexplored in the GI tract: Fe^3+^-mediated dopamine oxidation that forms *ortho*-quinones^43–45^ that cause α-syn to misfold and, subsequently, aggregate.^23, 24^ To this end, we again anaerobically cultured *E. coli* K-12 until stationary phase (14 hours) in either mM9_+NO3_ or mM9_-NO3_ but with the addition of α-syn monomer (20 µM) as well as dopamine (500 µM; mM9_+NO3,+DA_ or mM9_−NO3,+DA_, respectively) or its vehicle (mM9_+NO3,−DA_ or mM9_−NO3,−DA_, respectively). As before, bacteria cultured in nitrate-respiring conditions reduced nitrate to nitrite (Figure 3a). Accumulation of nitrite corresponded to an oxidizing redox potential (Figure 3b) and a shift in iron speciation so that the relative abundance of Fe^3+^ increased in comparison to cultures without nitrate supplementation (Figure 3c). Culture conditions in which nitrite was produced and dopamine was also present resulted in lower relative abundance of Fe^3+^ as compared to conditions without dopamine. This finding is putatively a reflection of dopamine oxidation being coupled to reduction of Fe^3+^, thereby increasing the relative abundance of Fe^2+^. Correspondingly, we were not able to measure redox potential in culture conditions that contained dopamine, as redox potential did not stabilize under these conditions.

**Figure 3.**
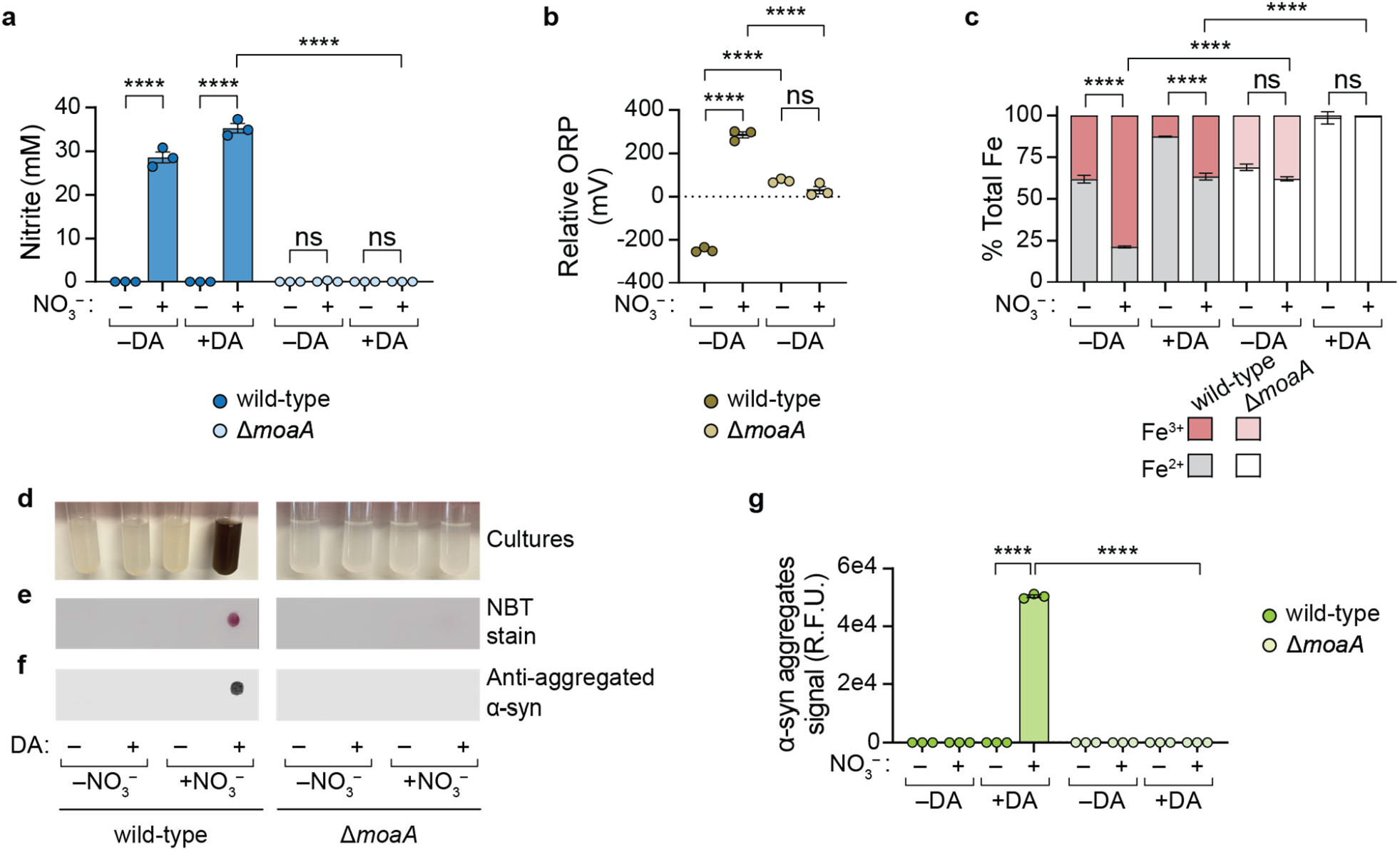
*E. coli* nitrate respiration instigates dopamine-dependent quinone formation and α-syn aggregation. *E. coli* K-12 wild-type or Δ*moaA* was cultured for 14 hours in mM9 media with α-syn monomer in the presence of nitrate (+NO_3_) or its absence (-NO_3_) and with dopamine (+DA) or without (-DA). Quantification of (a) nitrite (using Griess assay), (b) oxidation-reduction potential (ORP) relative to sterile media (using redox electrode), and (c) labile Fe^3+^ and Fe^2+^ (using ferrozine assay). (d–f) Representative images of (d) bacterial cultures as well as (e) membranes stained quinone formation (using nitroblue tetrazolium (NBT) stain) or (f) dot blots stained for α-syn aggregate formation (using immunostaining with anti-fibril α-synuclein as the primary antibody). (g) Quantification of α-syn aggregates in dot blots. R.F.U.: relative fluorescence units. n = 3 biological replicates; bars denote means ± S.E.M.; significance was determined using ordinary one-way ANOVA with Sidak’s multiple comparisons test; ****: *P* < 0.0001, ns: not significant.

In cultures containing dopamine, we observed formation of a dark pigment, which is characteristic of quinones (Figure 3d).^46^ Redox-cycling stain nitroblue tetrazolium (NBT), which specifically stains quinones,^47^ enabled detection of quinones in dot blots of nitrate-reducing bacterial cultures (Figure 3e and Figure S2). Contrastingly, and as depicted in Figure 3d, no dark pigment was observed in dopamine-supplemented fermentative culture conditions (mM9_−NO3,+DA_) wherein Fe^3+^ relative abundance was significantly less than in respiration culture conditions (mM9_+NO3,+DA_); correspondingly, no quinones were detected by NBT stain (Figure 3e; positive control shown in Figure S2). In the absence of dopamine supplementation to culture media, quinones were neither detected in nitrate-reducing nor in fermenting bacterial cultures. These data indicate that dopamine supplementation is necessary for quinone formation and that dopamine-dependent quinone formation occurs in the oxidizing conditions created upon bacterial reduction of nitrate.

Strikingly, dopamine-dependent quinone formation coincided with α-syn aggregation. Evaluation of α-syn aggregation was conducted by dot blot, immunostaining α-syn aggregates on membranes spotted with culture media (Figure 3f; positive control shown in Figure S3). Significantly greater amounts of α-syn aggregates formed in nitrate-reducing conditions as compared to fermentative conditions—but only when dopamine was present (Figure 3g and Figure S3). In the absence of dopamine, a culture environment of oxidizing redox potential (mM9_+NO3,−DA_; Figure 3b) was not sufficient to induce α-syn aggregation (Figure 3g and Figure S3).

We next used a genetic knockout of nitrate respiration to further clarify the roles of nitrate and nitrite in initiating α-syn aggregation. First, we targeted a molybdenum cofactor (MoaA) that is incorporated in the active site of nitrate reductases and is essential for this enzyme to reduce nitrate to nitrite.^48^ MoaA was deleted from *E. coli* K-12 wild-type to create the isogenic mutant *E. coli* K-12 Δ*moaA*. Culturing *E. coli* K-12 Δ*moaA* in mM9_+NO3_ media did not afford the growth advantage that was obtained when, upon culturing in the same media, *E. coli* K-12 wild-type performed nitrate reduction (Figure S4). Moreover, *E. coli* K-12 Δ*moaA* did not produce redox-active nitrite (Figure 3a), indicating that nitrate reduction was, indeed, inhibited by genetic deletion of *moaA*; likewise, a significantly less oxidizing redox potential was observed as compared to cultures of the wild-type strain cultured in mM9_+NO3_ media (Figure 3b). The less oxidizing redox potential of *E. coli* K-12 Δ*moaA* cultures corresponded to significant reductions in the relative abundance of Fe^3+^ when *E. coli* K-12 Δ*moaA* was cultured in either mM9_+NO3,−DA_ or mM9_+NO3,+DA_ as compared to analogous cultures of *E. coli* K-12 wild-type (Figure 3c). Without the ability of *E. coli* K-12 Δ*moaA* to reduce nitrate and increase the oxidizing redox potential of the culture media, neither dopamine oxidation nor α-syn aggregation occurred (Figures 3e, 3f, 3g; Figures S2 and S3). Taken together, these data indicate that the presence of nitrate, alone, does not induce α-syn aggregation; instead, we have demonstrated that bacteria that produce nitrite can transform an innocuous trio—Fe^2+^, dopamine, and α-syn monomers—into one that generates toxic α-syn aggregates.

### Tungstate inhibits α-syn aggregation induced by bacterial nitrate reduction

After identifying the instigating role of bacterial reduction of nitrate to nitrite in α-syn aggregation, we were curious about whether α-syn aggregation could be mitigated by chemically inhibiting bacterial nitrate respiration. To this end, we turned to tungstate, a chemical analog of molybdate that renders *Enterobacteriaceae* nitrate reductases inactive.^19^ We evaluated the effect of sodium tungstate (0.5–100 mM) on *E. coli* K-12 (wild-type and Δ*moaA*) cultured until stationary phase (14 hours) in mM9_+NO3_ media supplemented with α-syn monomer (20 µM) and with or without dopamine (Figure S5). For *E. coli* K-12 wild-type cultures, a dose-response relationship was observed: increasing concentrations of sodium tungstate resulted in decreasing production of nitrite (Figure 4a).

**Figure 4.**
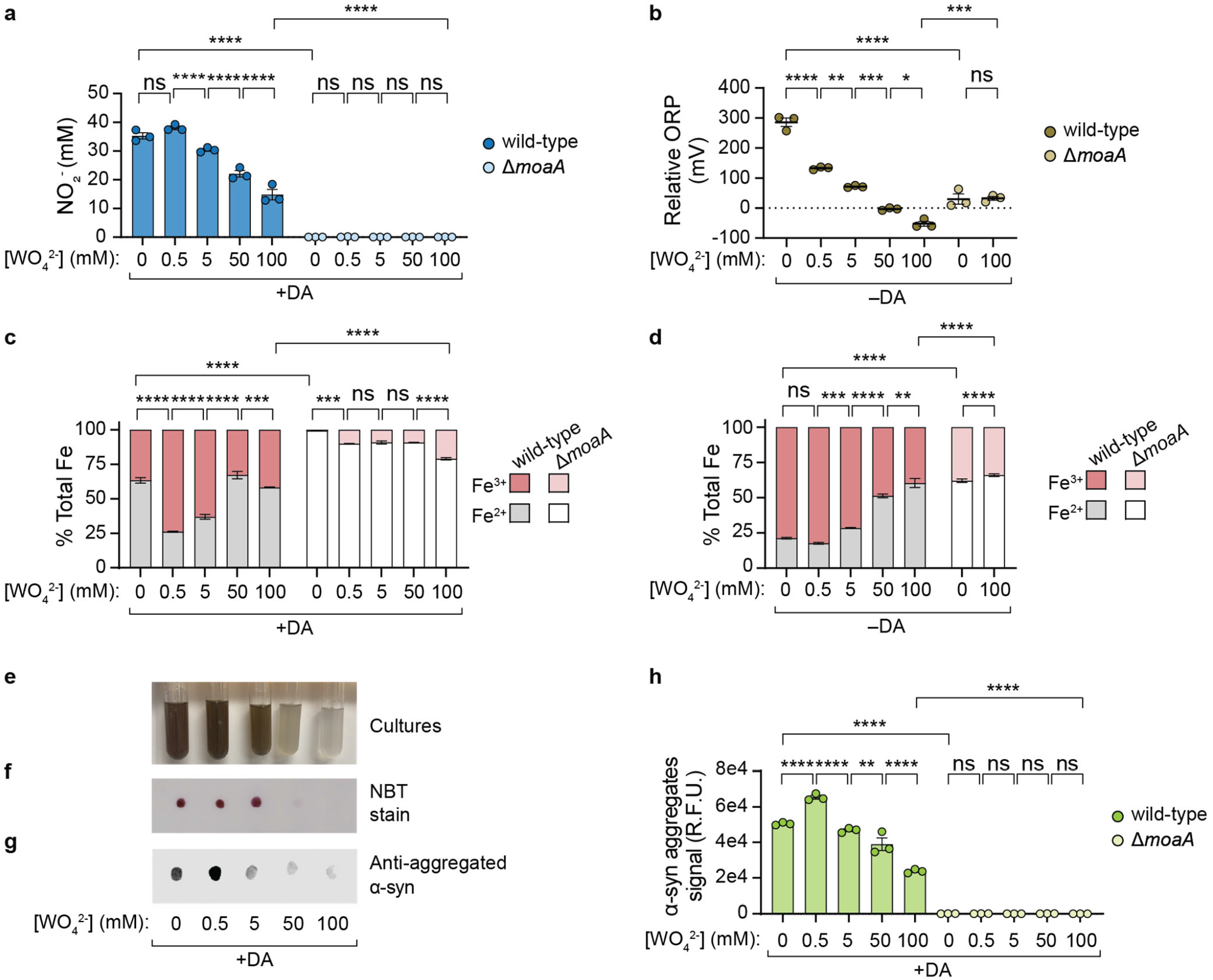
Tungstate limits α-syn aggregation by inhibiting bacterial nitrate reduction, which reduces redox potential of the bacterial environment. *E. coli* K-12 wild-type or Δ*moaA* was cultured for 14 hours in mM9 media with α-syn monomer in the presence of nitrate (+NO_3_), with dopamine (+DA) or without (-DA), and with sodium tungstate (WO_42-_) at varying concentrations (0–100 mM). Quantification of (a) nitrite (using Griess assay), (b) oxidation-reduction potential (ORP) relative to sterile media (using redox electrode), and (c–d) labile Fe^3+^ and Fe^2+^ (using ferrozine assay) in cultures (c) with dopamine or (d) without dopamine. (e–g) Representative images of (e) bacterial cultures of *E. coli* K-12 wild-type incubated in mM9_+NO3,+DA_ supplemented with tungstate (0–100 mM) as well as (f) membranes stained for quinone formation (using nitroblue tetrazolium (NBT) stain) or (g) dot blots stained for α-syn aggregate formation (using immunostaining with anti-fibril α-synuclein as the primary antibody). (h) Quantification of α-syn aggregates in dot blots. R.F.U.: relative fluorescence units. n = 3 biological replicates; bars denote means ± S.E.M.; significance was determined using ordinary one-way ANOVA with Sidak’s multiple comparisons test; ****: *P* < 0.0001, ***: *P* < 0.0007, **: *P* < 0.0026, *: *P* = 0.0168, ns: not significant.

In accordance with our findings that bacterial reduction of nitrate to nitrite creates a more oxidizing redox potential, progressive inhibition of this process using increasing concentrations of sodium tungstate supplemented to cultures of *E. coli* K-12 wild-type (in the absence of dopamine) correlated with decreasing oxidizing redox potentials of cultures (Figure 4b). As the redox potential of *E. coli* K-12 wild-type cultures decreased with increasing concentrations of tungstate, the relative abundance of Fe^3+^ also decreased in cultures with dopamine (Figure 4c) and without dopamine (Figure 4d). Tungstate (0.5–50 mM) inhibition of nitrate reduction by *E. coli* K-12 wild-type corresponded with decreased visible pigmentation of cultures (Figure 4e) and NBT-stained quinone (Figure 4f and Figure S2) as well as decreased formation of α-syn aggregates (Figures 4g and 4h; Figure S3). Although increasing tungstate concentrations from 0.5 mM to 100 mM resulted in significantly decreased nitrite level, relative redox potential, Fe^3+^ relative abundance, and α-syn aggregation, supplying *E. coli* K-12 wild-type cultures with up to 100 mM of tungstate did not ameliorate the effects of nitrite reduction to the extent that was observed upon genetic deletion of *moaA*. With 100 mM tungstate, nitrite concentration, Fe^3+^ relative abundance, and α-syn aggregates remained significantly greater in cultures of *E. coli* K-12 wild-type as compared with *E. coli* K-12 Δ*moaA*. Taken together, these results indicate that tungstate is a means to chemically limit (but not fully prevent) generation of the oxidizing environment created by *E. coli* nitrate reduction and, thereby, inhibit the cascade of oxidation reactions that lead to α-syn aggregation.

### Redox-active nitrite induces α-syn aggregation in specialized gut epithelial cells

We next set out to examine the relevance of our proposed bacteria-induced α-syn aggregation mechanism to the mammalian gut. In the GI tract, α-syn is expressed by specialized epithelial cells called enteroendocrine cells (EECs);^4^ the dopamine metabolic pathway is also expressed by these cells.^49^ EECs are chemosensory cells at the interface between gut luminal contents and the nervous system. While the apical side of these cells is in direct contact with the gut microbiome and its metabolites, a cellular projection (called a neuropod) on the basolateral surface of EECs forms synapses with enteric neurons, including those of the vagus nerve.^50^ Thus, EECs have been proposed as a potential site where environmental factors, including bacterial metabolites, could initiate α-syn misfolding and the prion-like cascade leading to PD.^4, 51^

Owing to gut epithelial cells absorbing nitrite through passive diffusion,^52^ we hypothesized that this redox-active metabolite that is produced in the gut lumen by nitrate-respiring bacteria^18, 19^ could induce aggregation of α-syn that is present in the cytoplasm of EECs.^4^ To test our hypothesis, we used murine STC-1 cells, an accepted model cell line for elucidating properties of native EECs.^53^ STC-1 cells were incubated with nitrate or nitrite (0.5–50 mM), and α-syn aggregation was analyzed via immunofluorescent staining using an antibody for α-syn fibrils (Figure 5a). STC-1 cells treated with 0.5 mM nitrate as compared to untreated cells showed no significant difference in amounts of α-syn aggregates (Figure 5b). In contrast, treatment with nitrite significantly induced α-syn aggregation in a concentration dependent manner, with 0.5 mM, 5 mM, and 50 mM nitrite resulting in 5.0-, 9.3-, and 10.8-fold increases in aggregation, respectively, as compared to untreated cells (Figure 5b). Additionally, at each concentration tested, aggregation was significantly elevated in nitrite-versus nitrate-treated cells. Taken together, these results not only provide strong evidence for our proposed model that nitrite induces α-syn aggregation within intestinal cells expressing this protein, but these findings also emphasize the importance of bacterial reduction of nitrate to nitrite to incite this pathogenic process.

**Figure 5.**
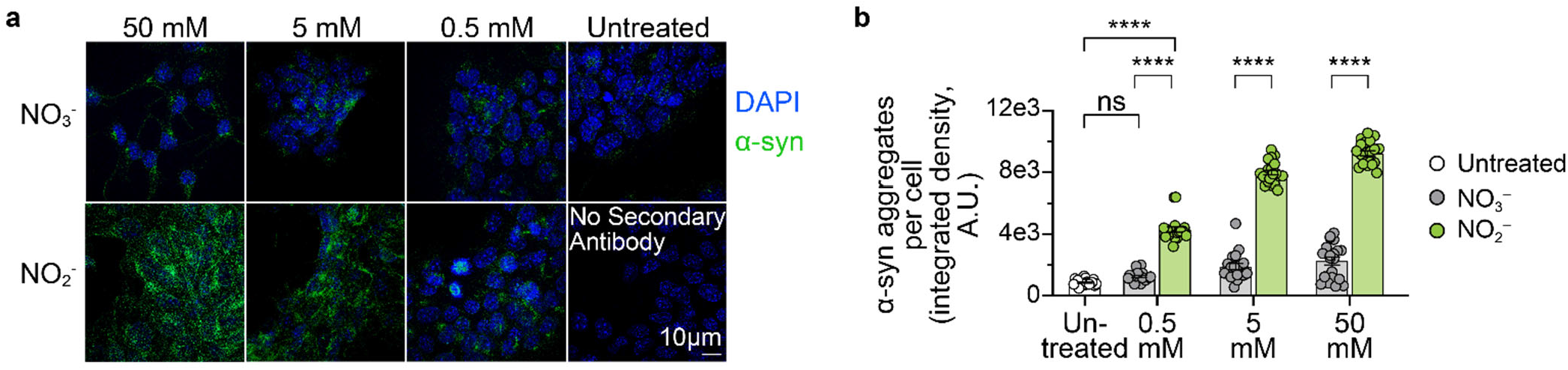
Nitrite, but not nitrate, induces α-syn aggregation in enteroendocrine STC-1 cells. α-syn aggregates per STC-1 cell upon incubation with nitrate (NO_3-_) or nitrite (NO_2-_) were (a) visualized (representative images) and (b) quantified using maximum intensity projections acquired by structured illumination microscopy (immunofluorescence staining of α-syn aggregates is in green; DAPI-stained cell nuclei are in blue). n = 20 cells; bars denote mean ± S.E.M.; significance determined by one-way ANOVA with Sidak’s multiple comparisons test; ****: *P <* 0.0001, ns: not significant.

### Nitrate-reducing bacteria provoke increased dopaminergic neurodegeneration in *Caenorhabditis elegans*

Having discovered that nitrate-reducing *E. coli* K-12 wild-type can initiate a cascade of reactions that results in α-syn aggregation *in vitro*, we next examined whether gut bacteria could modulate dopamine-dependent α-syn aggregation and neurodegeneration *in vivo*. For this investigation, we made use of the multiple features of *Caenorhabditis elegans* that position this organism as a valuable gnotobiotic model for discovery of molecular-level mechanisms governing the gut–brain axis.^54^ In particular, we selected transgenic *C. elegans* models of PD that have been used to elucidate dopamine’s role in the formation of toxic α-syn oligomers and in neurodegeneration.^55, 56^

First, we used *C. elegans* strain UA287 in which the six frontal dopaminergic neurons in this animal both express human α-syn A53T mutant and fluoresce by virtue of GFP, with expression of both proteins controlled by dopamine transporter promoter *P_dat-1_*.^55^ Neurodegeneration in these nematodes is evidenced by decline of the fluorescent signal (indicating neuron death) as well as by changes in the morphology of neuritic processes and soma (indicating decline in neuronal activity).^57, 58^ Previous studies using these animals have demonstrated that dopamine as well as oxidative damage that results from aging^55^ or environmental factors^59^ are mediators of α-syn-dependent dopaminergic neurodegeneration; yet, the influence of host–microbe interactions on this pathogenic process has remained undetermined.

To address this knowledge gap, *C. elegans* UA287 (n=15 hermaphroditic nematodes from each of three independent transgenic worm lines) were reared with a food source of either *E. coli* K-12 wild-type or Δ*moaA* aerobically cultured in mM9_−NO3_, both conditions that provided a baseline for neurodegeneration in the absence of bacterial nitrate reduction (Figure S6). Then, *C. elegans* UA287 synchronized L4 larvae (48 hours post-hatching) were exposed, in anaerobic conditions, to *E. coli* K-12 wild-type or Δ*moaA*, respectively, that had been anaerobically cultured in mM9_−NO3_, mM9_+NO3_, or mM9 supplemented with nitrite (mM9_+NO2_); after three hours, nematodes were returned to aerobic conditions, washed of bacterial treatments, and then resupplied with the respective food sources provided at baseline conditions. Following this acute exposure, neuronal decline was monitored using fluorescence microscopy every 48 hours until day 6 post-hatching, a time at which aging-independent neurodegeneration is typically observed.^60^ Next, dopaminergic neurodegeneration was assessed as previously reported^55, 58, 61^: nematodes were scored as having a neurodegenerative phenotype if any degenerative processes (e.g., a missing dendritic process, cell body loss, or a blebbing neuronal process) were observed.

To assess the effect of hypoxia on neurodegeneration, one group of nematodes was maintained in aerobic conditions, reared on a food source of *E. coli* K-12 wild-type (aerobically cultured in mM9_−NO3_), and not subjected to the acute bacterial exposure protocol (i.e., untreated controls). At days 4 and 6 post-hatching, 60±7% and 44±6%, respectively, of the untreated nematode population exhibited no dopaminergic neurodegeneration (Figures 6a and 6b). These observed rates of decline are consistent with previously reported rates of α-syn-dependent neurodegeneration observed in this *C. elegans* strain.^55^ Additionally, acute exposure of *C. elegans* UA287 to either *E. coli* K-12 wild-type or Δ*moaA* cultured in mM9_−NO3_ resulted in no significant differences in dopaminergic neurodegeneration as compared to untreated controls at day 4 or day 6 (Figures 6a and 6b). These data indicate that hypoxic conditions that nematodes were subjected to during the acute bacterial exposure protocol does not, itself, exacerbate neurodegeneration nor does bacterial genetic background, in the absence of nitrate, impact baseline levels of neurodegeneration.

**Figure 6.**
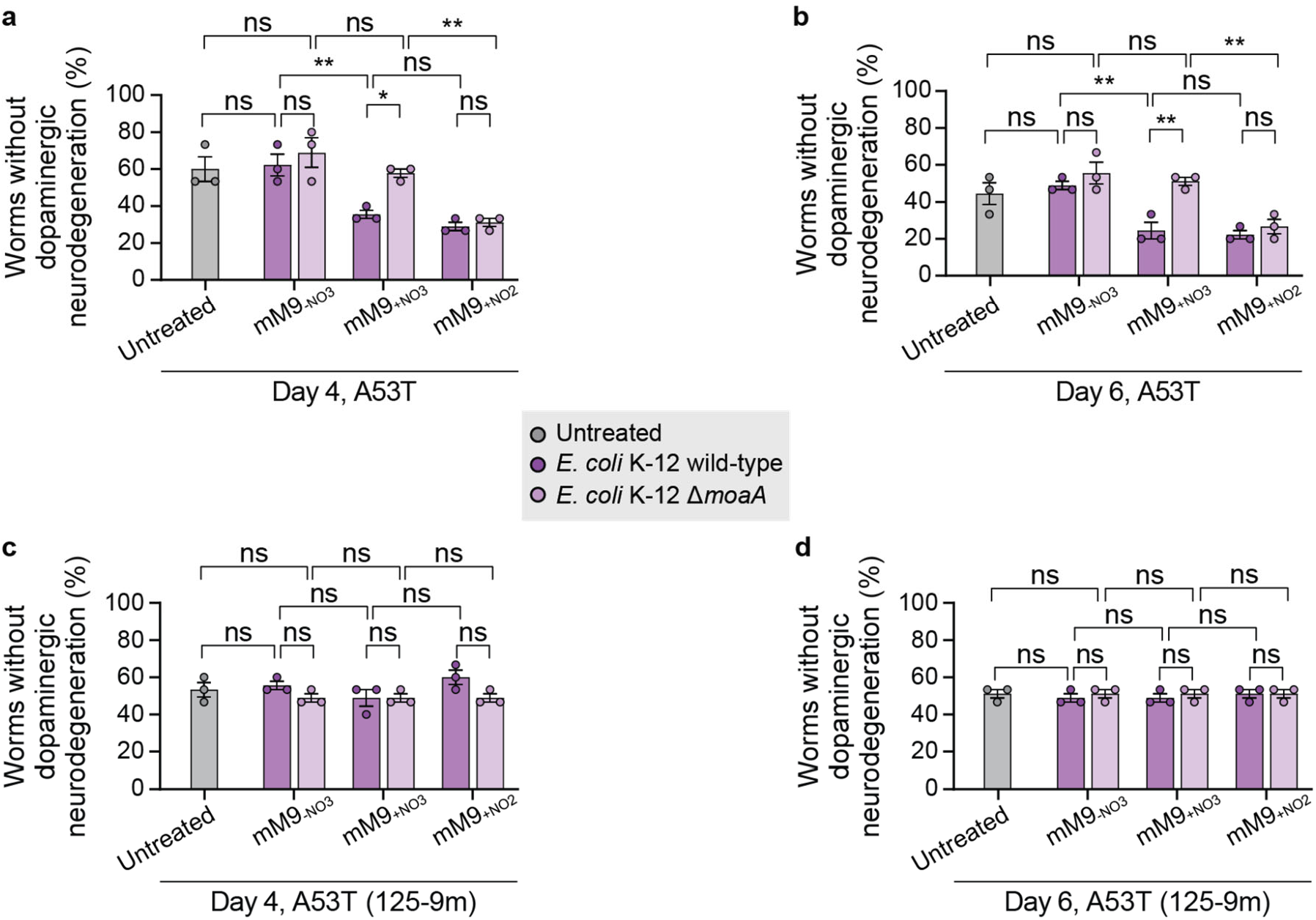
*E. coli* K-12 nitrate reduction induces dopaminergic neurodegeneration in *C. elegans* models of PD. Assessment of dopaminergic neurodegenerative phenotypes in *C. elegans* (a–b) strain UA287 (which expresses human α-syn A53T) and (c–d) strain UA288 (which expresses human α-syn A53T(125-9m)) following acute (three-hour) exposure to *E. coli* K-12 wild-type or Δ*moaA* cultured in mM9 media without nitrate (mM9_−NO3_) or mM9 media supplemented with nitrate (mM9_+NO3_) or nitrite (mM9_+NO2_). Acute bacterial exposure was performed under anaerobic conditions at 48 hours post-hatching. Neurodegenerative phenotypes were detected using fluorescence microscopy at (a,c) day 4 and (b,d) day 6 post-hatching. n = 45 nematodes; values are averages of 15 worms from each of three independent transgenic worm lines per genetic background; bars denote means ± S.E.M.; significance determined by one-way ANOVA with Sidak’s multiple comparisons test; *: *P* = 0.0102, **: *P* < 0.0039, ns: not significant.

Neurodegeneration phenotypes significantly increased when *C. elegans* UA287 were acutely exposed to cultures of *E. coli* K-12 wild-type in mM9_+NO3_. By day 4 post-hatching, only 36±2% of worms exhibited normal neurons (Figure 6a), which further dropped to 24±4% by day 6 (Figure 6b). Nitrate, alone, did not aggravate neurodegeneration in *C. elegans* UA287, as nematodes exposed to cultures of *E. coli* K-12 Δ*moaA* in mM9_+NO3_ displayed no significant difference in neurodegeneration as compared to untreated controls or to animals treated with *E. coli* K-12 Δ*moaA* cultured in mM9_−NO3_. In contrast, treating nematodes with *E. coli* K-12 wild-type or Δ*moaA* cultured with nitrite (mM9_+NO2_) resulted in significant neurodegeneration, which did not differ from levels observed upon treatment with cultures of *E. coli* K-12 wild-type in mM9_+NO3_. As worms aged from day 4 to day 6, there were fewer worms without neurodegenerative phenotypes across every treatment group while the relative trends in neurodegeneration were maintained across both time points (Figures 6a and 6b). Taken together, these findings indicate that *C. elegans* UA287 exposed to nitrate-reducing *E. coli* K-12 does not merely induce dopaminergic neurodegeneration but also that a single acute exposure to the products of this bacterial metabolic pathway—nitrite—is enough to induce persistent and progressive neuronal decline.

### Interaction between dopamine and α-syn are critical determinants of whether nitrate-reducing bacteria induce dopaminergic neurodegeneration in *C. elegans* model of PD

Owing to data from our *in vitro* experiments that showed the necessity of dopamine to α-syn aggregation that was induced by bacterial nitrate reduction, we next sought to determine the significance of dopamine to this mechanism of neurodegeneration *in vivo*. To examine this mechanism, we used *C. elegans* strain UA288.^62, 63^ This strain has the same genetic background as strain UA287, except that a mutated version of α-syn is encoded: five amino acids (residues 125–129) near the C-terminus of α-syn were mutated from Y_125_EMPS_129_ to F_125_AAFA_129_ [mutant referred to as A53T(125-9m)]. These five mutations prevent the interaction between dopamine quinones and the C-terminus of α-syn that leads to α-syn aggregate formation.^62, 63^

When dopamine was prevented from interacting with α-syn, bacterial culture conditions that induced dopamine-dependent quinone formation and α-syn aggregation *in vitro* as well as neurodegenerative phenotypes in *C. elegans* UA287 had no impact on dopaminergic neurodegeneration in *C. elegans* UA288. Neither acute exposure of *C. elegans* UA288 to *E. coli* K-12 wild-type cultured in either mM9_+NO3_ or in mM9_+NO2_ nor *E. coli* K-12 Δ*moaA* cultured in mM9_+NO2_ had any impact on neurodegeneration as compared to untreated nematodes at either day 4 (Figure 6c) or day 6 (Figure 6d). Mirroring findings from our *in vitro* experiments, these data indicate that nitrite, alone, is not sufficient to induce a neurodegenerative phenotype. Instead, nitrite drives dopamine-dependent α-syn aggregation as a mechanism of neurotoxicity in this *C. elegans* model of PD.

## DISCUSSION

Here, we show that the ability of *Enterobacteriaceae*, specifically *E. coli*, to modulate the redox potential of a bacterium’s environment plays a critical role in inducing the formation of α-syn aggregates. Microbial metabolism has been previously demonstrated to influence the redox potential of the environment;^34^ however, gut bacterial metabolic pathways are typically associated with creating more reducing environments, while the generation of oxidizing environments is commonly linked to host processes.^29–33, 64^ Our data suggest that bacteria performing nitrate dissimilatory metabolism can generate an oxidizing environment. When this metabolic process occurs, *Enterobacteriaceae* reduce nitrate, a relatively redox-inert by-product of the host inflammatory response,^65^ to nitrite, an oxidizing agent.

Using *E. coli* K-12 wild-type cultures, we demonstrated that the presence of nitrate in the growth medium results in its reduction to nitrite as well as a generation of a redox environment that is more oxidizing as compared to the same cultures without nitrate. In contrast, nitrate supplied to bacteria with a nitrate respiration defect (i.e., *E. coli* K-12 Δ*moaA*) or to sterile media neither resulted in nitrite production nor a more oxidizing redox potential. These data indicate that *E. coli* K-12 nitrate metabolism generates an oxidizing environment. Notably, we showed that the shift in redox potential that accompanies nitrite production was sufficient to alter the relative abundance of iron species so that Fe^3+^ predominated over Fe^2+^ in cultures. This shift occurred in spite of anaerobic and reducing *in vitro* culture conditions similar to those of the GI tract that favor the prevalence of Fe^2+^ over Fe^3+^.^39^ *Enterobacteriaceae* nitrate respiration may be an underappreciated mechanism by which gut bacteria disrupt their environment’s relative abundance of Fe^2+^ and Fe^3+^ as well as the metabolic processes mediated by this redox-active metal.

Iron has been implicated in PD onset due to the ability of Fe^3+^ to oxidize dopamine to *ortho*- quinones that cause α-syn monomers to aggregate.^24, 66, 67^ Through *in vitro* experiments, we demonstrated that gut bacterial nitrate reduction can induce the cascade of oxidation reactions that ultimately results in dopamine-dependent α-syn aggregation. This is the first report elucidating a gut bacterial metabolic pathway that directly influences α-syn aggregation *in vitro*. If conserved in the mammalian GI tract, this biochemical pathway may be a novel target for intervention strategies to prevent α-syn aggregation in the gut.

Due to tungstate’s inhibition of *Enterobacteriaceae* nitrate respiration,^19^ we sought to determine whether tungstate could be used to inhibit *Enterobacteriaceae*-induced α-syn aggregation. Tungstate exposure limited dopamine oxidation and α-syn aggregation in cultures of *E. coli* K-12 wild-type supplemented with nitrate. Additionally, tungstate treatment effectively lessened the oxidizing redox potential of the bacterial environment as well as increased the relative abundance of less-oxidizing Fe^2+^. Owing to the ability of oral tungstate treatment to effectively ameliorate murine colitis, which is exacerbated by gut bacterial nitrate respiration,^19^ the ability of tungstate to limit α-syn aggregation *in vitro* may have important therapeutic implications for limiting α-syn aggregation in the mammalian intestine.

Towards determining the significance of bacterial nitrate respiration to α-syn aggregation in the gut, we focused our efforts on EECs. EECs are emerging as a critical mediator of the gut–brain axis^68, 69^ and have been implicated in PD as a source of intestinal α-syn.^4, 51^ Since EECs can form synapses with the enteric nervous system,^68^ it has been hypothesized that α-syn aggregates may spread from EECs to the brain via the vagus nerve;^4, 51^ however, the precise molecular stimuli of α-syn aggregation in EECs have remained elusive. Owing to our findings that nitrite induces dopamine-dependent α-syn aggregation *in vitro*, we suspected that nitrite could induce the same process in EECs. Within EECs, nitrite exposure (0.5–50 mM) afforded a dose-dependent increase in the amount of α-syn aggregation. Consistent with our *in vitro* experiments, the effect of nitrate (0.5 mM) on α-syn aggregation was no different than sham treatment. Notably, the concentration of nitrate in the mucus layer of the intestinal lumen is on the order of 0.5 mM,^18^ which supports the physiological relevance of our findings. We demonstrated that nitrite— the product of gut bacterial *Enterobacteriaceae* nitrate respiration that occurs within the GI tract^18, 19^—can induce α-syn aggregation in EECs, which line the GI lumen.

*Enterobacteriaceae* is more abundant in people with PD as compared to non-diseased, age-matched controls and is positively correlated with the severity of motor dysfunction;^9^ however, whether *Enterobacteriaceae* plays a causative role in PD has remained unknown. Findings from our *in vitro* experiments that demonstrated the capacity of nitrite to induce α-syn aggregation provided the foundation to test a causative role for bacterial nitrate respiration on α-syn-dependent neurodegeneration *in vivo* using a *C. elegans* model of PD. Our results demonstrated that acute exposure of nematodes to *E. coli* K-12 wild-type that reduced nitrate to nitrite caused significant acceleration of dopaminergic neurodegeneration as compared to either treatment with cultures of *E. coli* K-12 wild-type without nitrate or to no bacterial treatment. While genetic knockout of nitrate respiration (Δ*moaA*) in *E. coli* K-12 precluded this bacterium from inducing neurodegeneration, even in the presence of nitrate, exposing nematodes to the redox-active metabolic product of nitrate reduction, nitrite, recapitulated the dopaminergic neurodegenerative phenotype that was evoked by nitrate-reducing *E coli* K-12. In addition to the significance of nitrite, the neurodegenerative phenotype depended on the presence of α-syn’s C-terminus residues (125–129) that interact with dopamine quinones to instigate α-syn aggregate formation.^62, 63^ Mutation of these α-syn residues [A53T(125-9m)] to preclude interactions between α-syn and dopamine quinone abrogated neurodegeneration: neither exposure to nitrate-reducing bacteria nor nitrite could induce neurotoxicity beyond the levels observed in untreated controls. These findings suggest that acute exposure of *C. elegans* to the metabolic product of bacterial nitrate reduction induces the putative formation of dopamine-derived quinones that, upon interaction with the C-terminus of α-syn, causes neurotoxic α-syn aggregation.

In summary, our data demonstrate that the gut microbiota, in particular *E. coli* (a prototypic organism of the *Enterobacteriaceae* bacterial family^18^), is capable of inducing α-syn aggregation *in vitro* and α-syn-associated neurodegeneration in a *C. elegans* model of PD. Here, we identified a specific metabolic pathway, bacterial nitrate reduction, that can generate the oxidative environment that causes dopamine oxidation and subsequent α-syn aggregation. While dopamine oxidation has been identified as a crucial component of α-syn aggregation mechanisms in the brain,^24, 66, 67^ we have demonstrated that dopamine-dependent mechanisms of α-syn aggregation are also likely relevant in the gut. Our findings that nitrite induced α-syn aggregation in EECs provides strong motivation for examining these cells as a likely conduit of the gut–brain axis in PD. Future work will focus on the ability of α-syn aggregates to spread to the enteric nervous system following their formation in EECs. Finally, our studies of *C. elegans* models of PD provide supporting evidence for the *in vivo* relevance of gut bacteria nitrate reduction to α-syn aggregation and neurodegeneration. If the cascade of reactions initiated by bacterial nitrate reduction and ending with α-syn aggregation is conserved in the mammalian gut, our findings position gut bacterial nitrate reduction as a novel target that may be leveraged for early intervention strategies to prevent intestinal α-syn aggregation and limit Parkinsonian neurodegeneration.

## MATERIALS AND METHODS

Full details for all materials and methods are provided in the Supplementary Materials.

## ASSOCIATED CONTENT

**Supplementary Materials:** Materials and Methods; Figures S1-S6

## AUTHOR INFORMATION

**Corresponding Author:** Elizabeth N. Bess, Departments of Chemistry and Molecular Biology & Biochemistry, University of California, Irvine, California, USA.

**Author Contributions:** L.O.O. and E.N.B. developed the project and performed data analysis. L.O.O. designed and performed experiments with contributions from K.S.U. L.O.O., K.S.U., and E.N.B provided critical feedback on experiments. L.O.O. and E.N.B. wrote the manuscript. E.N.B. acquired funds and provided project supervision and administration. All authors read and approved the final version of the manuscript.

**Notes:** The authors declare no conflicts of interest. All authors read and approved the final version of the manuscript. Data generated or analyzed during this study are included in the manuscript and supporting files or are available from the corresponding author upon reasonable request.

## ACKNOWLEDGMENTS

We thank Professor Kimberlee Caldwell and Professor Guy Caldwell for kindly providing the *C. elegans* strains used in this study. The table of contents graphic was created with BioRender.com (agreement number DM240BYT0B). This work was supported by the University of California, Irvine School of Physical Sciences and the University of California Cancer Research Coordinating Committee (C21CR2124). This study was made possible in part through access to the Optical Biology Core Facility of the Developmental Biology Center, a shared resource supported by the Cancer Center Support Grant (CA-62203) and Center for Complex Biological Systems Support Grant (GM-076516) at the University of California, Irvine.

