## Supplementary Material for "Discovery of a Gut Bacterial Metabolic Pathway that Drives α-Synuclein Aggregation and Neurodegeneration"

### Material and Methods

Bacteria and culture conditions

*General*

*Escherichia coli* K-12 BW25113 was acquired from the Coli Genetic Stock Center. *Escherichia coli* Δ*moaA* was constructed as previously described.[1,2] Archived stocks of bacteria were maintained in 16% glycerol at -80 °C. Bacteria were routinely cultured in modified M9 mineral media (mM9, defined below) at 37 °C under anoxic and reducing conditions (2–5% $H_2$, 20% $CO_2$, with the balance being $N_2$) in a COY anaerobic chamber. When indicated, mM9 media (described below) was supplemented with 50 mM sodium nitrate (Sigma, S5506-250G), 50 mM sodium nitrite (Fisher Scientific, M1065490100), 500 µM dopamine HCl (Alfa Aesar, A11136-06) and/or 20 µM α-synuclein (purified as described below). Prior to beginning experiments, media was equilibrated in anoxic conditions overnight to remove oxygen.

mM9 media was prepared to contain the following: 1x M9 salts (Sigma, M6030-1KG), 2 mM magnesium sulfate (Fisher Scientific, 0338-500G), 100 µM calcium chloride (Fisher Scientific, AC219171000), 20 mM D-glucose anhydrous (VWR Life Science, Biotechnology Grade), 0.2% (*w/v*) casamino acids (Fisher Scientific, DF0288-15-6), 1x vitamin supplement (ATCC, MD-VS), and 500 µM ferrous iron chloride (Oakwood, 098678-5g).

*Purification of recombinant α-synuclein*

Purification of α-synuclein was performed according to the method of Huang et al., 2005.[3] Briefly, an overnight culture of *E. coli* Rosetta 2 transformed with pET-21a-α-synuclein (Addgene, 51485) was diluted 100-fold in LB supplemented with 100 µg/mL ampicillin. Expression of α-synuclein was induced for 5 hours by adding 100 µM IPTG to the culture media when cell density reached an $OD_{600}$ of 0.3–0.4. The cell pellet from a 1 L culture was resuspended in 100 mL of osmotic shock buffer (30 mM Tris–HCl, 40% sucrose, and 2 mM ethylenediaminetetraacetic acid disodium, pH 7.2) and incubated for 10 minutes at room temperature. The pellet collected by centrifugation at 12,000 rpm for 20 min was quickly resuspended in 90 mL of cold water followed by adding 37.5 µL of saturated $MgCl_2$. The resuspension was kept on ice for 3 min. The supernatant containing periplasm proteins was collected by centrifugation at 3500 x *g* for 20 min and dialyzed overnight against buffer A (20 mM Tris–HCl, pH 8.0). After centrifugation at 3500 x *g* for 20 min, the supernatant was loaded onto a DEAE Sepharose column (GE healthcare) and eluted with a 0–0.5 M NaCl gradient in buffer A. The elution fractions were analyzed by SDS–15% PAGE, and the fractions containing only an 18 kDa band were combined for size-exclusion chromatography in 0.0095% $MgCl_2$, 0.316% Tris–HCl, 0.58% NaCl, pH 7.5. Fractions containing pure α-synuclein were concentrated to 1 mg/mL, aliquoted, and stored at -20 °C.

*Redox potential measurements*

Redox potential was measured in 5 mL bacterial cultures using a redox electrode with an Ag/AgCl reference electrode (Cole-Parmer, EW-59001-75). The values are reported relative to the standard hydrogen electrode (SHE) and were determined by measuring the offset of the reference electrode in a reference solution with a known redox potential (Thermo Scientific, 967961). Relative oxidation-reduction potential (ORP) was determined by subtracting the ORP value of the sterile control from the ORP of experimental conditions.

*Nitrate and nitrite measurements*

To determine nitrate concentrations, the Griess-vanadium chloride method developed by Miranda *et al.* was used, with slight modifications.[4] Briefly, 2 µL of bacterial culture was diluted in 198 µL of 1 M HCl. Of this suspension, 50 µL were mixed with 50 µL of a solution containing 0.1 % (*w/v*) sulfanilamide (Sigma, S9251-100G) and 0.05% (*w/v*) *N*-(1-naphthyl)ethylenediamine dihydrochloride (Sigma, N9125-10G) in 0.5 M HCl followed by the rapid addition of 50 µL of 0.25 % (*w/v*) vanadium (III) chloride (Sigma, 208272-1G) in 1 M HCl. The mixture was incubated at 37 °C for 30 minutes. Nitrite was evaluated in a similar manner except that samples were exposed to 50 µL of 1 M HCl instead of vanadium (III) chloride. In either case, following incubation the absorbance at 540 nm was measured using a SPECTROstar Nano plate reader (BMG LABTECH). Sample concentrations were determined using calibration curves generated by measuring absorbance at 540 nm and performing linear regression against known concentrations of nitrate and nitrite prepared from sodium nitrate or sodium nitrite standards, respectively, at concentrations ranging from 1000 to 15.62 µM (2-fold dilution series).

*Iron speciation assay*

To determine the ratio of $Fe^{2+}/Fe^{3+}$ in media, we used the $Fe^{2+}$-ferrozine assay as previously described, with slight modifications.[5] Briefly, to determine the concentration of $Fe^{2+}$ present in media, bacterial culture (70 µL) was mixed with 70 µL of 50 mg/mL ferrozine (Hach, 230424) in 1 M potassium acetate buffer (pH 5.5). To determine total iron present in the solution, ascorbic acid was used as a reducing agent to reduce $Fe^{3+}$ to $Fe^{2+}$. For this purpose, bacterial culture (90 µL) was mixed with 100 µL of 50 mg/mL ferrozine in 1 M potassium acetate buffer (pH 5.5) and 10 µL of 1 M ascorbic acid (Sigma, A92902-25G). Reduced and non-reduced samples were incubated at 37 °C for 30 minutes followed by centrifugation at 3000 x *g* for 15 minutes. Then, 100 µL of supernatant was used to determine the light absorption of $Fe^{2+}$-ferrozine complex at 562 nm using a SPECTROstar Nano plate reader (BMG LABTECH). Sample concentrations were determined using calibration curves generated by measuring absorbance at 562 nm and performing linear regression against known concentrations of $Fe^{2+}$ prepared from $FeCl_2$ standards at concentrations ranging from 100 to 1.56 mM (2-fold dilution series). $Fe^{3+}$ was calculated as the difference between total iron (the sum of $Fe^{2+}$ and $Fe^{3+}$ reduced to $Fe^{2+}$) and $Fe^{2+}$.

*Nitroblue tetrazolium assay for the detection of quinones*

The presence of quinones in bacterial cultures was evaluated as previously described.[6] Briefly, under ambient conditions, 2 µL of each sample was spotted onto a polyvinylidene difluoride (PVDF) membrane (Fisher Scientific, ISEQ00005) that was previously activated in methanol and equilibrated in tris-buffered saline (pH 7.4). As a positive control, 2 µL of oxidized dopamine produced by *Aspergillus* tyrosinase (Worthington Biochemical, LS003789; detailed protocol described below) was used. The membrane was left to dry for at least one hour. Then, the membrane was reactivated using methanol and submerged in a solution of 0.6 mg/mL nitroblue tetrazolium (Santa Cruz Biotechnology, sc-296003) in a potassium glycinate buffer (pH 10). The membrane was then incubated at room temperature, in the dark, for 45 minutes. After incubation, the membrane was washed twice in a solution of 0.16 M sodium borate (Sigma, S9640-25G) and photographed using an Epson V19 scanner.

*Dopamine oxidation using tyrosinase*

Lyophilized *Aspergillus* tyrosinase (Worthington Biochemical, LS003789) was resuspended in 1000 µL 50 mM phosphate buffer, 15% glycerol, pH 6.5, to create a 24 kU/mL solution (50 mg/mL, 390 µM). Single-use aliquots (100 µL) were prepared and stored at -20 °C. Aliquots were thawed on ice shortly before use. Dopamine (500 µM) was oxidized by adding 250 U (3.9 µM) of tyrosinase. The solution was incubated for one hour at room temperature.

*Immunodetection of α-synuclein aggregates*

α-Syn aggregation was evaluated in methanol-activated PVDF membranes. Under anaerobic conditions, 2 µL of bacterial cultures were spotted and left to dry for at least one hour. As controls, 2 µL of monomeric (Abcam, ab51189) and aggregated α-synuclein (Abcam, ab218817) were each spotted into the membrane. The membrane was immunostained with anti-fibril α-synuclein (Abcam, ab209538; 1:50,000 dilution) as primary antibody and NIR 800CW donkey anti-rabbit IgG secondary antibody (LI-COR, 926-32213) according to the manufacturer's instructions. The membranes were visualized using an Odyssey CLx imager (LI-COR) using the

800 nm channel. α-syn aggregates were quantified using Image studio data analysis software (LI-COR).

Cell lines and growth conditions

*General*

Enteroendocrine STC-1 (CRL-3254) cell line was obtained from the American Type Culture Collection (ATCC). STC-1 cells were routinely cultured in Dulbecco's Modified Eagle Medium, containing 25 mM glucose and 1 mM sodium pyruvate (DMEM; Thermo Fisher Scientific, Gibco 11995065), supplemented with 10% fetal bovine serum (FBS; Thermo Fisher Scientific, Gibco 10-437-010), and incubated at 37 °C in an humidified atmosphere containing 5% $CO_2$. Cells were seeded onto two 8-well glass slides (Stellar Scientific, NST230104) at a density of $1x10^5$ viable cells and incubated for 48 hours. Then, the growth media was replaced with fresh media supplemented with either sodium nitrate, sodium nitrite, or vehicle. Cells were incubated for another 24 hours before fixation.

*Immunofluorescence*

Fixation was performed using 10% formalin (Fisher Scientific, 22-170-402) for 20 minutes at room temperature. Cells were then permeabilized for another 20 minutes at room temperature using 0.1% Triton X-100 (Bio-Rad, 1610407) in phosphate-buffered saline (PBS), pH 7.4. After discarding the permeabilization solution, wells were washed with PBS, twice. Samples were then blocked for 1 hour at room temperature in PBS containing 5% Normal Goat Serum (Thermo Scientific, 50197Z) and 0.2% bovine serum albumin (BSA; Fisher Scientific, 501613336) followed by washing three times using PBS. Subsequently, the glass slide chamber was incubated at 4 °C overnight with anti-α-syn aggregate primary antibodies (Abcam, ab209538; dilution 1:250). The chambers were then washed with PBS followed by an incubation with anti-goat Alexa Fluor-488 secondary antibody (Abcam, ab150077; dilution 1:500) for 1 hour at room temperature in the dark.

Finally, the glass slides were mounted using 8 drops of Invitrogen ProLong Diamond Antifade Mountant with DAPI (Fisher Scientific, P36962). A 24 mm x 60 mm coverslip (VWR, 48393-106) was carefully placed on top, avoiding the formation of air bubbles. The slides were cured for 24 hours in the dark.

*Structured illumination microscopy*

Images were acquired under identical conditions using a Zeiss Elyra 7 super-resolution microscope with a x63 oil immersion lens. Images were collected using 405 nm and 488 nm laser lines for excitation; emission filters used were BP 420-480 (DAPI) and BP 495-525 (Alexa fluor 488). Z-stack images with an interval of 1 μm were acquired and deconvoluted with the 3D-SIM algorithm using the strong default reconstitution settings. The maximum projection of the z-stack was then obtained using Zen Black 3.0 software. Sections were imaged in the absence of primary antibodies, and images were captured at the same gain as images with primary antibody. No endogenous tissue fluorescence was observed in the absence of primary antibodies. Quantification of the fluorescence signal was performed using Image J software; an area of 100 $\mu m^2$ around each nucleus was used to facilitate segmentation of the cells. For each condition, 20 cells were evaluated. Results are expressed as arbitrary units (AU) of fluorescence intensity measured using the integrated density option.

<u>Nematode strains and growth conditions</u>

*General*

*C. elegans* strains were handled and maintained at 20 °C according to standard practices.[7,8] The following strains were used: UA287 (P*dat-1*::A53T α-synuclein, P*unc-54*::tdTomato];*vtIs7* [P*dat-1*::GFP) and UA288 (P*dat-1*::A53T(125-9m) α-synuclein, P*unc-54*::tdTomato];*vtIs7* [P*dat-1*::GFP).[9] Worms were synchronized using the alkaline hypochlorite method[8] and left nutating overnight at room temperature to hatch in sterile M9 buffer (3 g/L

$KH_2PO_4$ [Fisher Scientific, P285500], 6 g/L $Na_2HPO_4$ [Sigma, S3264-250G], 5 g/L NaCl [Sigma, S7653-250G], 1 mM $MgSO_4$). Synchronized germ-free worms were routinely grown using Nematode Growth Media (NGM)[8] supplemented with *E. coli* K-12. Three independent transgenic worm lines were analyzed per genetic background.

*Acute exposure to anaerobic bacterial cultures*

Synchronized L1 nematodes were reared, aerobically, in NGM liquid media supplemented with 6 mg/mL of a food source of either *E. coli* K-12 wild-type or Δ*moaA* aerobically cultured in $mM9_{-NO3}$ until stage L4, providing baseline conditions for neurodegeneration in the absence of bacterial nitrate reduction. Next, synchronized L4 larvae (48 hours post-hatching) were centrifuged at 280 x *g* for two minutes and then washed five times with M9 buffer to remove the bacterial diet. Then, approximately 5,000 worms (as calculated in 10 µL droplets as previously reported[8]) were exposed for three hours, under anaerobic conditions, to 5 mL of *E. coli* K-12 wild-type or *E. coli* K-12 Δ*moaA*. Conversely, untreated worms were resupplied with *E. coli* K-12 wild-type (aerobically cultured in $mM9_{-NO3}$). Anaerobic bacterial cultures were obtained by culturing in mM9 media (described above) for exactly 14 hours at 37 °C under anaerobic conditions; mM9 was supplemented with 50 mM sodium nitrate, 50 mM sodium nitrite, or vehicle, as specified.

After three hours of exposure, the nematodes were returned to aerobic conditions, washed five times in M9 buffer, then resupplied with the food source provided at baseline conditions, and finally plated on NGM plates seeded with the *E. coli* food source that had been provided at baseline conditions. NGM plates were also supplemented with 120 µM 5-fluoro-2-2deoxyuridine (FUdR; VWR, IC10555110), an inhibitor of DNA synthesis that blocks egg-hatching. Until day 6 post-hatching, nematodes were transferred to new plates every 48 hours; at the time of each transfer, neuronal decline was monitored using fluorescence microscopy, as detailed below.

*Analysis of neurodegeneration*

Dopaminergic neurodegeneration was analyzed as previously described.[9–11] Briefly, 15 hermaphroditic nematodes were randomly selected to examine each nematode's six frontal dopaminergic neurons using fluorescence microscopy (Zeiss Elyra 7 super-resolution microscope with a x48 oil immersion lens). Nematodes were scored as having a neurodegenerative phenotype if any degenerative processes (e.g., a missing dendritic process, cell body loss, or a blebbing neuronal process) were observed. Percentages of worms without neurodegenerative phenotypes were reported. Three independent transgenic worm lines were analyzed per genetic background.

Data and statistical analysis

Data were plotted and statistically analyzed using GraphPad Prism 8 software. Unless stated otherwise, means ± standard error of the mean (S.E.M.) were plotted. Statistical tests performed are specified in figure legends. For all analyses, $P \leq 0.05$ was considered significant. Sample sizes are noted within the main text, figure legends, and within Materials and Methods.

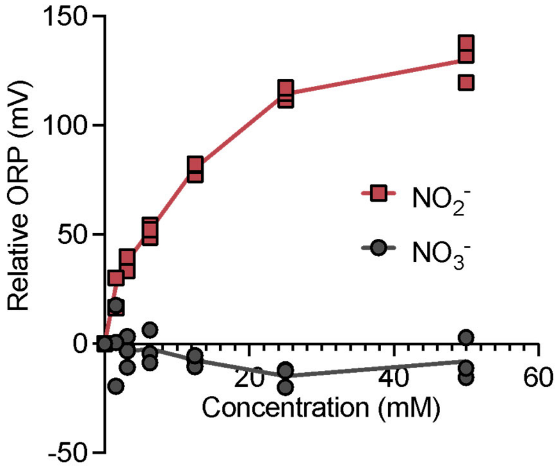


**Figure S1.** Effect of sodium nitrate and sodium nitrite on redox potential upon supplementation to sterile mM9 media. n=3 technical replicates.

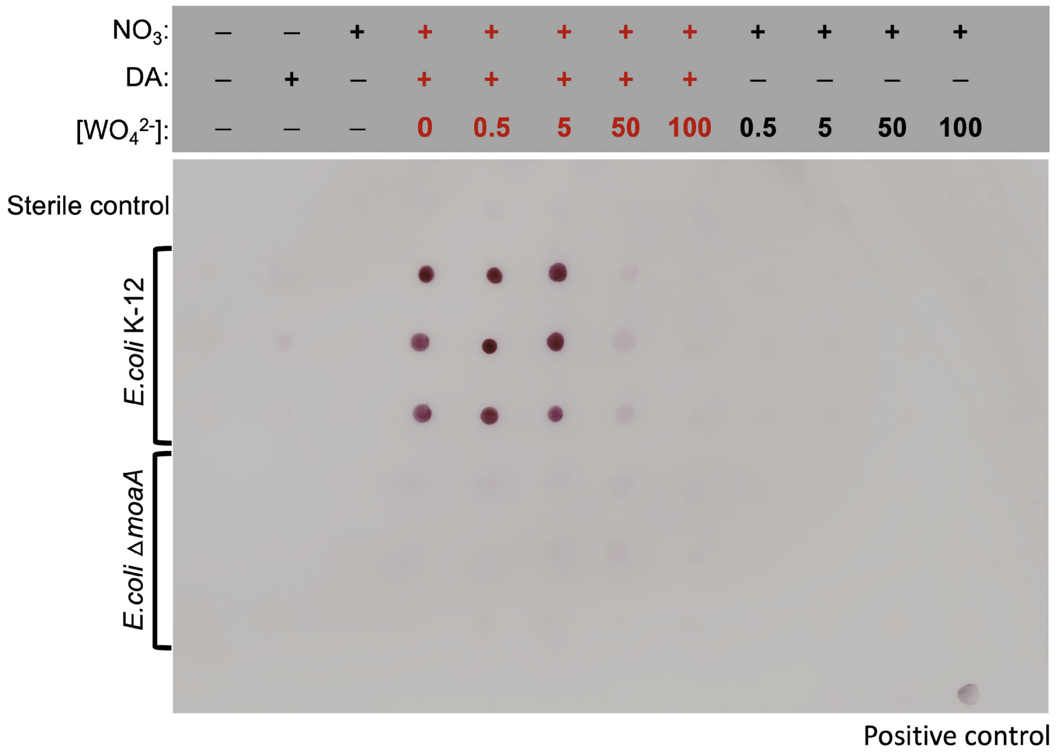


**Figure S2.** Detection of quinones by NBT staining of bacterial cultures spotted into PVDF membranes. The positive control was dopamine that was oxidized using *Aspergillus* tyrosinase and subsequently spotted on the membrane.

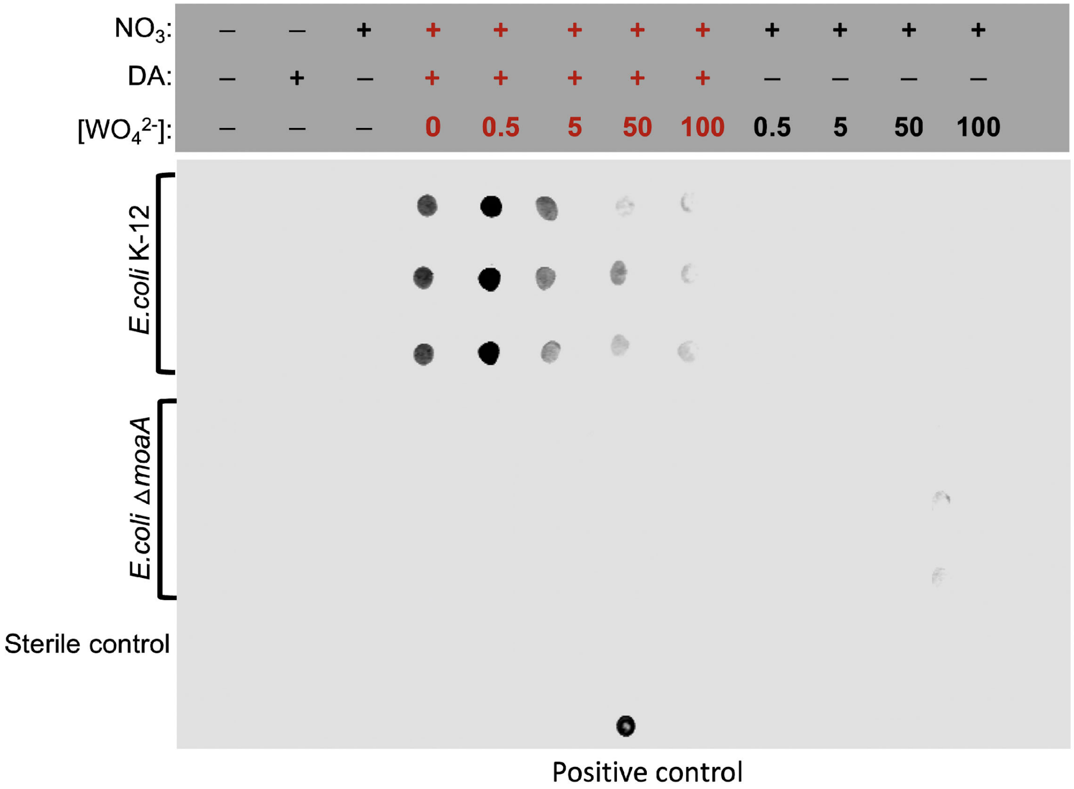


**Figure S3.** Detection of α-synuclein aggregation by immunostaining using anti-fibril α-synuclein as primary antibody. The positive control was commercial aggregated α-synuclein.

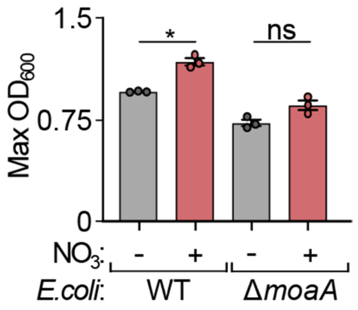


**Figure S4.** Maximum optical density at 600 nm ($OD_{600}$) of *E. coli* K-12 wild-type or Δ*moaA* cultures after 14 hours of incubation in the presence or absence of nitrate supplementation to mM9 culture media. n = 3 biological replicates; bars denote means ± S.E.M; significance was determined using one-way ANOVA with Turkey's multiple comparisons; *: *P*=0.0278; ns: not significant.

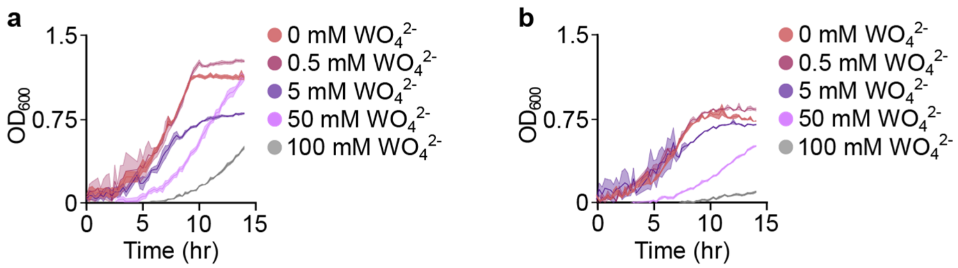


**Figure S5.** Effect of increasing concentrations of sodium tungstate on growth of bacterial cultures of (a) *E. coli* K-12 wild-type and (b) *E. coli* K-12 Δ*moaA* incubated for 14 hours in mM9$_{+NO3}$.

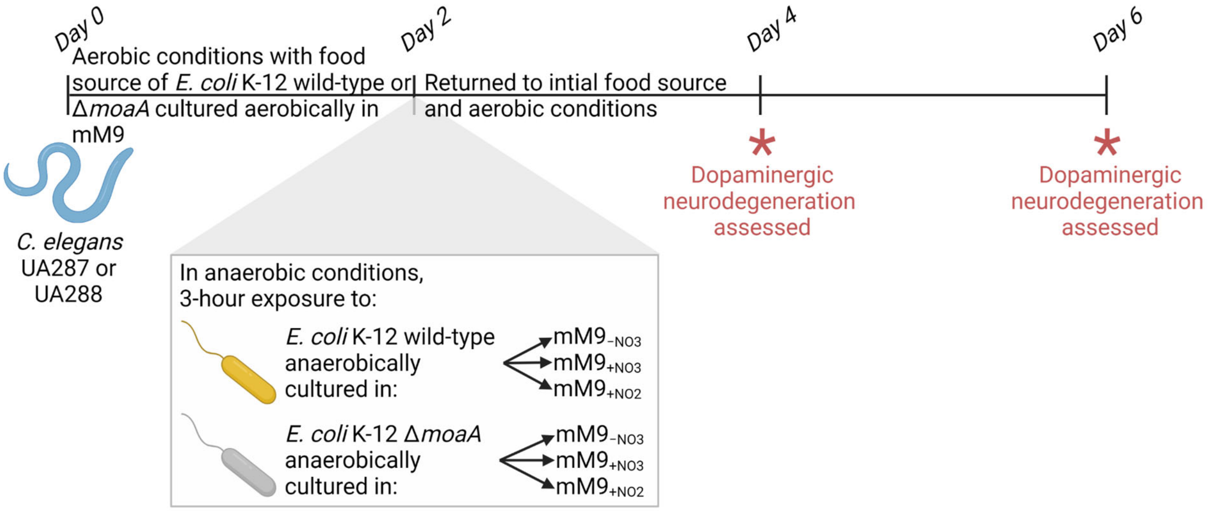


**Figure S6.** Schematic of experiments performed using *C. elegans* UA287 and UA288. (Figure created with BioRender.com, agreement number HT240BYHXD.)
